# The effects of morning priming exercise on afternoon physical and cognitive performance in male rugby union players

**DOI:** 10.1101/2025.05.29.656928

**Authors:** Jamie Knight, Mark Russell, Daniel Cunningham, Christian Cook, Mark Waldron, Liam Kilduff

## Abstract

**Objective:** To assess the effect of morning resistance exercise on afternoon physical and cognitive performance in male rugby union players.

**Methods:** On two occasions (randomised, crossover design), 25 male rugby union players completed morning physical (countermovement jump, 40 m linear sprint) and cognitive (rapid visual information processing; spatial working memory; paired associates of learning) assessments. Control (passive rest) or intervention (barbell back squat, 3 x 3 repetitions at 85% of one repetition maximum and barbell squat jump, 5 x 3 repetitions at 40% one repetition maximum) trials were implemented 5.5 h before afternoon assessments.

**Results:** Differences between morning and afternoon assessments were found in sprinting performance following intervention for 30 m fly (-0.08 ± 0.13 s, *P =* 0.001, ES: -0.75) and 40 m (-0.12 ± 0.17 s, *P =* 0.004, ES: -0.62) relative to control (0.03 ± 0.07 s, 0.01 ± 0.09 s, 30 m fly and 40 m, respectively). Afternoon countermovement jump flight time improved following intervention compared to control (2% vs. 0%, *p =* 0.012). Delta analysis discovered significant differences from morning to afternoon in intervention (0.385 ms ± 0.053) but not control (0.405 ms ± 0.057). No changes were found for cognitive assessments (*P* > 0.05).

**Conclusion:** Morning lower body resistance exercise, incorporating a combination of high intensity and ballistic exercises, was an effective morning strategy to improve afternoon markers of physical but not cognitive performance in male rugby union players.

## Introduction

Match outcomes in rugby union (RU) can be influenced by several key performance indicators, incorporating both physical and technical-tactical components [1]. Effective ball carrying, for example, has been associated with successful match outcome and is partially underpinned by physical qualities, such as lower body peak power output and acceleration momentum [1, 2, 3]. Additionally, rugby players must constantly adapt to changing situations and make situation specific decisions under extreme pressure and fatigue, highlighting the importance of the cognitive demands within the sport [4]. A window of opportunity exists on competition day to acutely enhance physical performance [5], with morning-based resistance training (RT) implemented five to six hours prior to competition being efficacious in improving indices of afternoon neuromuscular performance [6, 7, 8, 9].

Rugby union requires multiple cognitive skills, such as attention and working memory (WM) [10]. Previous researchers have demonstrated that athletes with a high WM capacity are better able at focusing their attention on tactical decision-making whilst blocking out irrelevant auditory distractions [11]. Likewise, Scharfen and Memmert [12] reported WM capacity to be positively related to sport specific skills within elite youth footballers. Indeed, resistance exercise has been shown to acutely enhance both attention and WM [13, 14], although the effects of morning RT priming on these cognitive skills remains unclear. While not all research agrees [7], some evidence exists to suggest that cognitive performance can be improved by morning RT [15]. Professional cricketers have displayed improved Stroop test performance (-3.83 s, 7.4%) compared to control following completion of morning RT [15]. Conversely, Russell et al. [7] reported no significant improvements in reaction time following morning RT in male RU players. Mechanistically, testosterone has been positively linked to brain function [16]. It is prudent to note that resistance exercise can acutely elevate testosterone concentrations and it is feasible that attenuating such declines may protect indices of performance later in the day [6, 7, 17].

From a neuromuscular perspective, improvements in countermovement-jump (CMJ) peak power (+2.7%) and 40 m sprints (+1.3%) have been reported six hours after morning high intensity RT [6]. More recently, morning lower body RT (i.e., trap bar deadlifts, 6 x 4 repetitions up to 85% repetition maximum; RM) enhanced afternoon sprinting performance (-0.07 s, 1.2%) and jump height (+1.1 cm, 2.5%) within professional male cricketers [15]. Several mechanisms, such as increased mechanical stiffness, motor neuron activation, and increased fibre sensitivity to calcium ions have been proposed to explain the efficacy of RT for enhanced afternoon performance [8]. Likewise, diurnal changes in hormonal status have been suggested to play a substantial role [6].

Low-volume, and high-load (≥85% 1RM) traditional exercise or low-load (30-40% 1RM) ballistic exercise have been recommended as most effective in a recent review [8]. Additionally, it has been proposed that movement-specific activity be performed within priming sessions as these appear to be more effective in comparison to general, non-specific activity [9]. A notion supported by previous performance improvements in jumping, throwing, and sprinting when the priming activity implemented a similar movement pattern [6, 18, 19]. Contrary to this, Russell et al [7] prescribed upper body RT (bench press 5 x 10 at 75% RM) in the morning to professional rugby players and noted significant improvements in the first two repetitions of a repeated sprint protocol. Demonstrating that movement-specific priming activity may not be more beneficial than general movement, however, further research is required to elucidate these findings. This may be more beneficial for RU players due to the wide range of movements required to be successful. Further research is needed to establish an effective priming strategy for RU players across positions.

The potential for morning RT to acutely improve indices of cognitive performance may be particularly beneficial for RU players on game day. Therefore, this study aimed to investigate the effect of a morning RT priming session on afternoon cognitive and physical performance in male RU players. We hypothesised that a morning RT priming session would influence afternoon physical and cognitive performance.

## Materials and methods

Following institutional ethical approval (Swansea University Ethics Committee; 3 2023 7344 6572), 25 amateur male rugby union players (age 21 ± 1 years, body mass 99.7 ± 12.5 kg, stature 1.86 ± 0.07 m) volunteered to participate in the current study following a recruitment period. All participants were healthy, injury-free and provided written informed consent prior to involvement.

Participants completed a control (CON) and intervention (INT) assessment day, each separated by four days in a randomised, counterbalanced, cross-over design. Participants were asked to maintain consistent nutritional practices and to abstain from caffeine, alcohol, strenuous exercise and any cognitively demanding tasks in the preceding 24 hours of each assessment day. Trial day timings (Fig. 1) remained consistent across both assessment days to avoid the influence of diurnal variation in performance [20]. Participants were familiarised with the testing battery seven days prior to the study commencing and had over 12 months RT and maximal sprint training experience.

**Fig 1.**
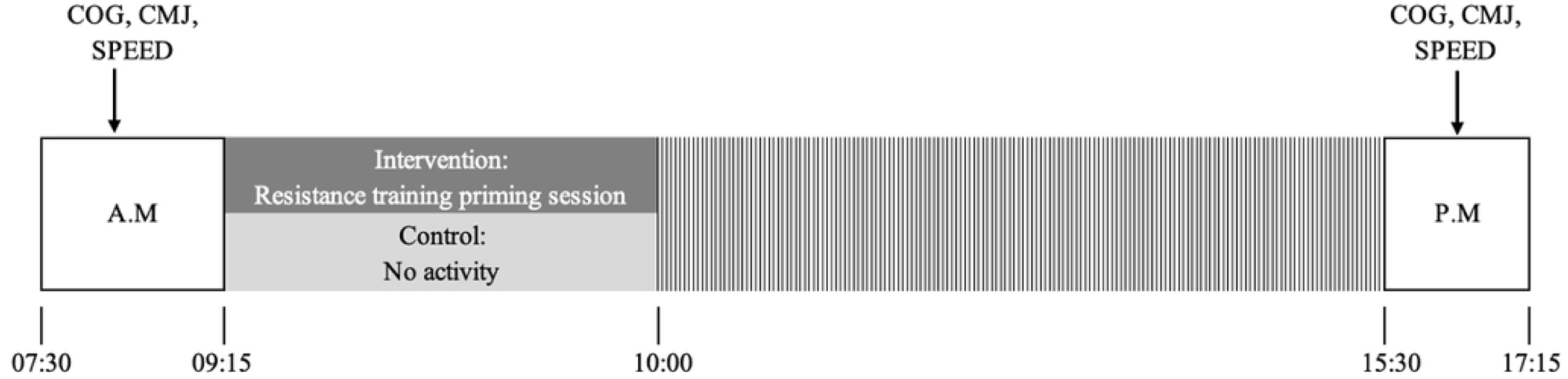
Overview of study protocol. COG = Paired associates of learning, spatial working memory, and rapid visual information processing assessments; CMJ = counter-movement jump; SPEED = 40 m linear sprint; 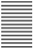 = passive recovery; A.M = morning base measures; P.M = afternoon measures.

Upon arrival at the testing facility, participants initially completed three cognitive assessments via the Cambridge Neuropsychological Test Automated Battery (Cambridge Cognition Ltd) (CANTAB). These were undertaken on an iPad in a quiet room with minimal distractions and have been found to be both valid and reliable assessments of cognitive ability [21]. The CANTAB test administration is fully automated, with participants being provided with on-screen text instructions and voiceover guidance for each assessment, explaining response requirements and task goals. The participants were sat at a comfortable distance from the iPad and were instructed to use the index finger of their dominant hand during the tests. Following these assessments, participants completed a standardised dynamic warm-up consisting of stretching and a 5-min moderate cycle followed by the countermovement jump (CMJ) assessment. Participants then completed a standardised sprint warm-up followed by a 40 m linear sprint assessment on an artificial grass pitch. The participants then underwent the priming intervention (INT) or the passive recovery period (CON) dependent on trial allocation. A 5.5-h passive recovery period followed for both INT and CON prior to participants completing afternoon testing using the same cognitive and physical testing battery in the same order as morning testing.

For INT, participants completed lower-body resistance exercise following a standardised warm-up (∼8 min) that required completion of dynamic stretching and progressive cycle (Table 1).

**Table 1.** Priming intervention completed by participants.

| Exercise | Sets | Reps | Intensity (% of 1<br>RM) | Rest<br>(minutes) |
| --- | --- | --- | --- | --- |
| Barbell Back Squat | 1 | 3 | 50 | 1 |
|  | 1 | 3 | 75 | 3 |
|  | 3 | 3 | 85 | 3 |
| Barbell Squat Jump | 5 | 3 | 40 | 3 |

*RM = repetition maximum*

### Cognitive performance assessment

The CANTAB assessments are widely used within various clinical populations and have demonstrated their validity and test-retest reliability with traditional neurocognitive tests [21, 22, 23, 24, 25]. The CANTAB assessments in the current study consisted of the Paired Associates Learning (PAL), the Rapid Visual Information Processing (RVP) and the Spatial Working Memory (SWM) assessment completed in this order at all assessment points.

### Paired associates learning

The PAL assessment was used to assess visual memory, new learning, and episodic memory [26]. The PAL is an 8-minute assessment where boxes are displayed on the screen and open one by one in a randomised order to reveal patterns hidden inside. The patterns are then displayed in the middle of the screen, one at a time. Participants were required to touch the box where the pattern was originally located. Patterns are re-presented to remind the participant of their locations if an error was made.

### Rapid visual information processing

The RVP assessment was used for the assessment of sustained attention [26]. The RVP is a 7- minute assessment where single digits appear one at a time at a rate of 100 digits per minute. Participants were required to detect a series of target sequences and touch a button when they identified the last digit of a target sequence.

### Spatial working memory

The SWM assessment was used to assess working memory and executive functioning [26]. A 4-minute assessment where the test begun with coloured boxes being shown on the screen. The aim of the test was that by touching the boxes and using a process of elimination, the participant should find one ‘token’ in each of the boxes and use them to fill up an empty column on the right-hand side of the screen. Participants were instructed that the computer will never hide a token in the same-coloured box, so once a token is found, the participant should not return to that box to look for another token. The colour and position of the boxes used are changed from trial to trial to discourage the use of stereotyped search strategies.

Physical performance was assessed via a CMJ and 40 m linear sprint. All jumps were performed on a Kistler force plate (model 9260AA, Kistler, Germany) and followed the previously recommended protocol for CMJ assessment [27]. Each CMJ commenced in a static standing position with arms akimbo, from which participants performed a preparatory dip before explosively jumping to attain maximum height. Participants performed three repetitions on each occasion, with the greatest flight time achieved being retained for analysis. The 40 m linear sprint assessment was conducted on an artificial grass pitch used for training, with participants wearing their normal footwear on both occasions. Sprints commenced 0.5 m behind the start line, where participants waited for the verbal start command. Time taken to complete the 40 m sprint, along with a 10 m split time was recorded using electronic timing gates (Brower Timing Systems, USA) with the fastest of three completed sprints being retained for analysis.

### Statistical analysis

Statistical analyses were carried out using SPSS Statistics software (IBM Inc., USA, version 29) with significance set at *P* ≤ 0.05 and data reported as mean ± standard deviation (SD). The Shapiro-Wilk test for normality was performed along with Mauchly’s test for sphericity with the Greenhouse-Geisser correction applied if the assumption of sphericity was violated. Two-way repeated measures ANOVA were performed to test for interaction effects (condition x time (A.M and P.M)). Where significant interaction effects were observed, trial was deemed to have influenced responses and simple main effect analyses were undertaken. Significant differences were investigated via post-hoc Bonferroni-adjusted pairwise comparisons and partial eta-squared (η^2^) values. Paired t-tests were completed to discover any differences between time points and trials where significant interaction effects were found. Hedge’s g effect sizes (ES) were calculated for post-hoc comparisons, and were interpreted as trivial (0.00-0.19), small (0.20-0.49), moderate (0.50-0.79), or large (≥ 0.80) [28].

## Results

### Sprint Performance

Fig 2a-c presents the sprint data as a function of trial. A significant interaction effect was found for both 30 m fly *(P* <0.001, partial-eta^2^ = 0.386) and 40 m sprints (*P* = 0.004, partial- eta^2^ = 0.302). Post-hoc analysis for A.M to P.M comparisons elicited superior mean changes in 30 m fly (-0.08 ± 0.13 s, *P =* 0.001, ES: -0.75, *moderate*) and 40 m (-0.12 ± 0.17 s, *P =* 0.004, ES: -0.62, *moderate*) sprint times following INT compared to CON (0.03 ± 0.07 s, 0.01 ± 0.09 s, 30 m and 40 m respectively). In addition, Table A in S1 File present the linear sprint data.

**Fig 2.**
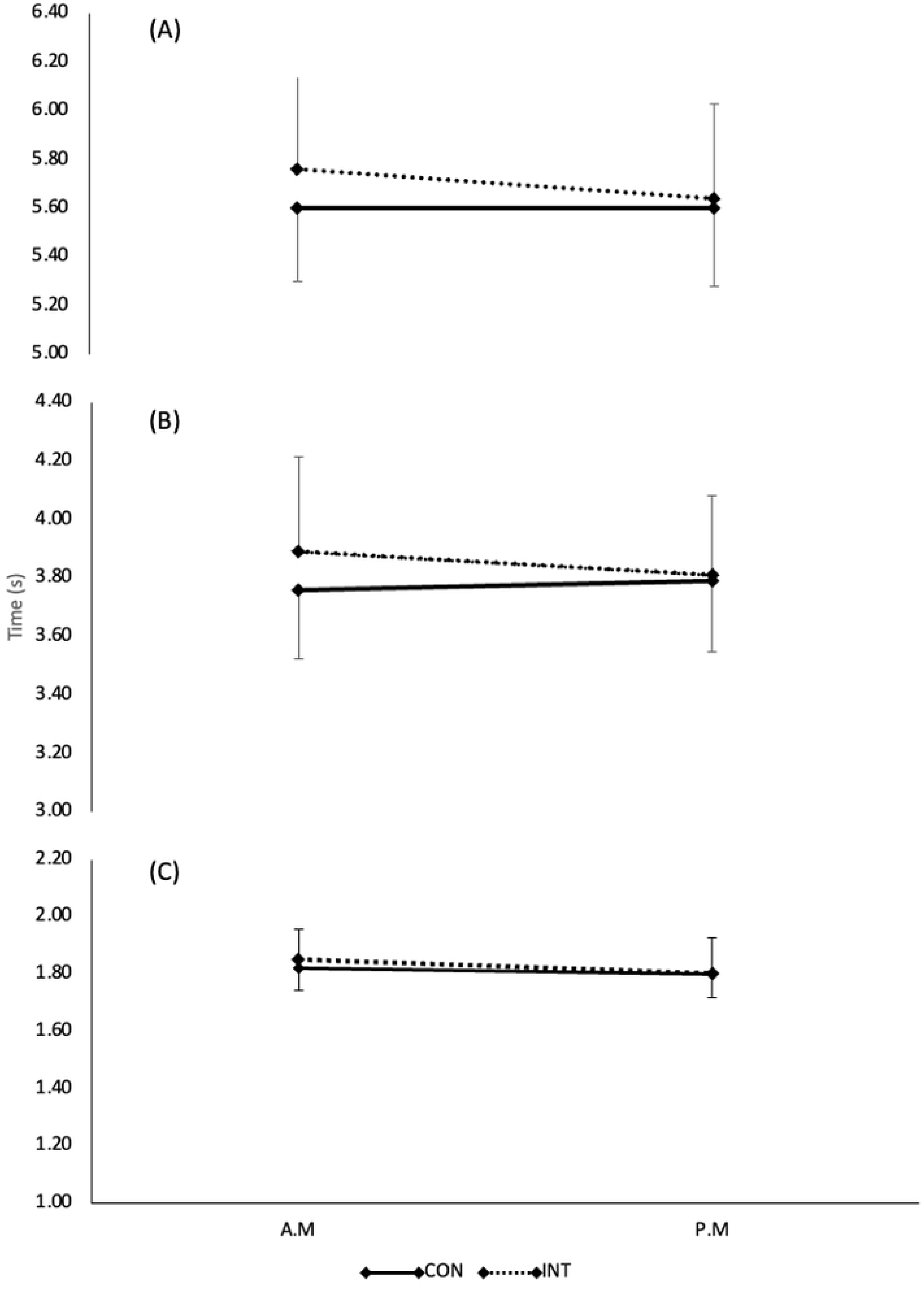
Mean ± SD linear sprint times (s) for (A) 40 m (B) 30 m fly (C) 10 m across conditions and time points.

### Countermovement Jump Performance

A significant interaction effect was found for CMJ FT (*P* = 0.012, partial-eta^2^ = 0.237) and was significantly influenced by time (*P* = 0.004, partial-eta^2^ = 0.291). Similar responses were observed between trials at A.M (0.402 ms ± 0.561; t_(24)_= 2.049, *p* = 0.052) and P.M (0.385 ms ± 0.053; t_(24)_= -0.494, *p* = 0.626), but P.M data in INT exceeded CON (2% vs. 0%; t_(24)_= -2.730, *p =* 0.012, ES: -0.53). Post hoc analysis discovered significant differences from A.M to P.M in INT (0.385 ms ± 0.053; t_(24)_= -3.628, *p* = 0.001, ES: -0.7) compared to CON (0.405 ms ± 0.057; t_(24)_=-0.10, *p* = 0.992) (Fig 3). Table B in S1 File present the CMJ test data.

**Fig 3.**
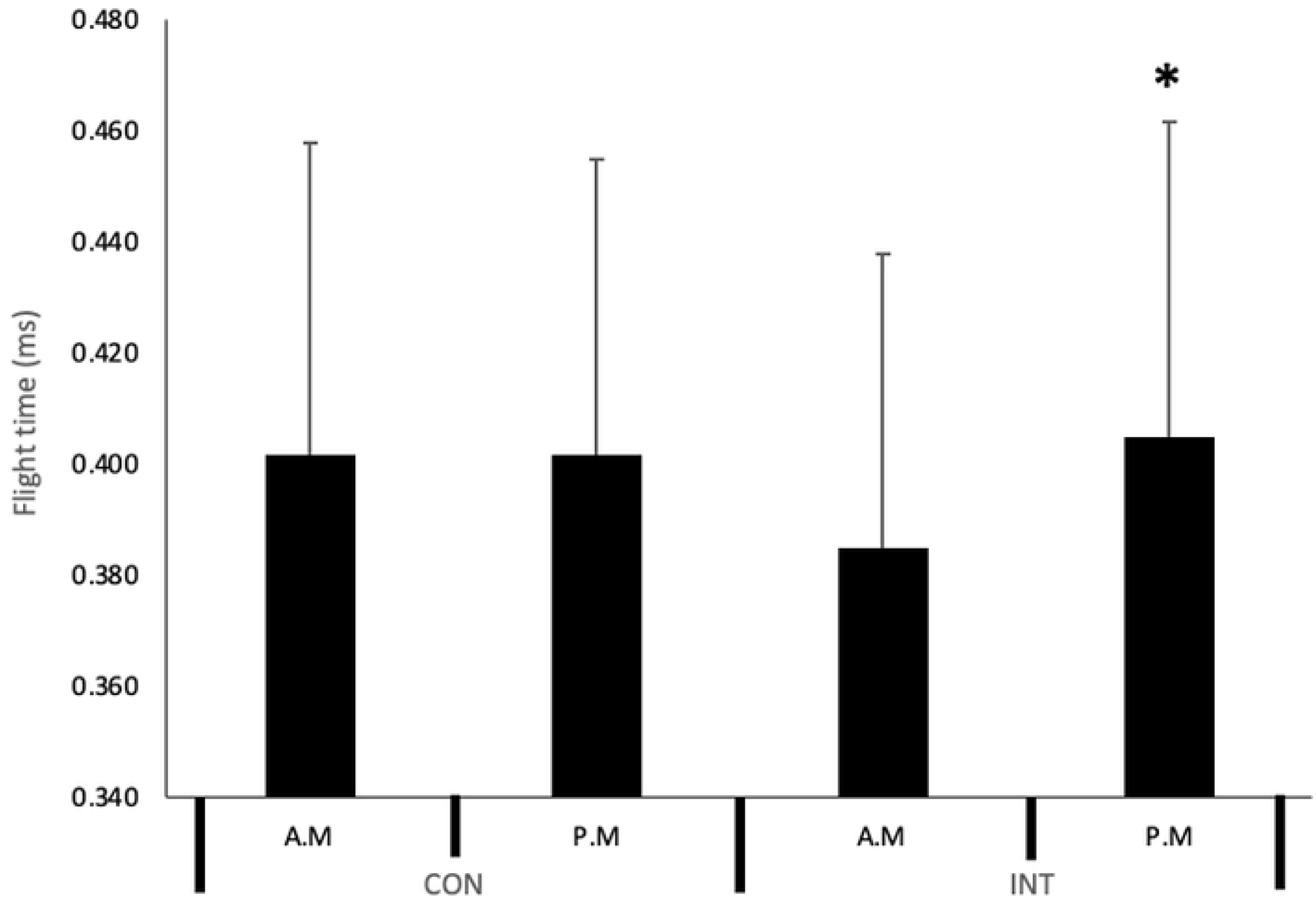
Mean ± SD CMJ flight time across conditions and time points. CON = Control, INT = Intervention. * = significant difference to A.M time point of trial condition.

### Cognitive assessments

No significant interaction effects were found for any indices of cognitive performance (PAL, RVP or SWM) (Table 2). A significant time effect was found for RVP mean response latency of a correct response to a target sequence (RVPML) (F_(1,24)_ = 5.710, *p* = 0.025, partial-eta^2^ = 0.192). Although non-significant, response latencies reduced from A.M to P.M in CON (453.53 ± 75.27 vs. 436. 07 ± 50.82, ES: 0.33, -3.75%) and INT (468. 24 ± 83.55 vs. 442.81 ± 52.82, ES: 0.34, -5.56%). No other significant time effects were found in the cognitive test outputs. In addition, Table C in S1 File presents the CANTAB test data.

**Table 2.**
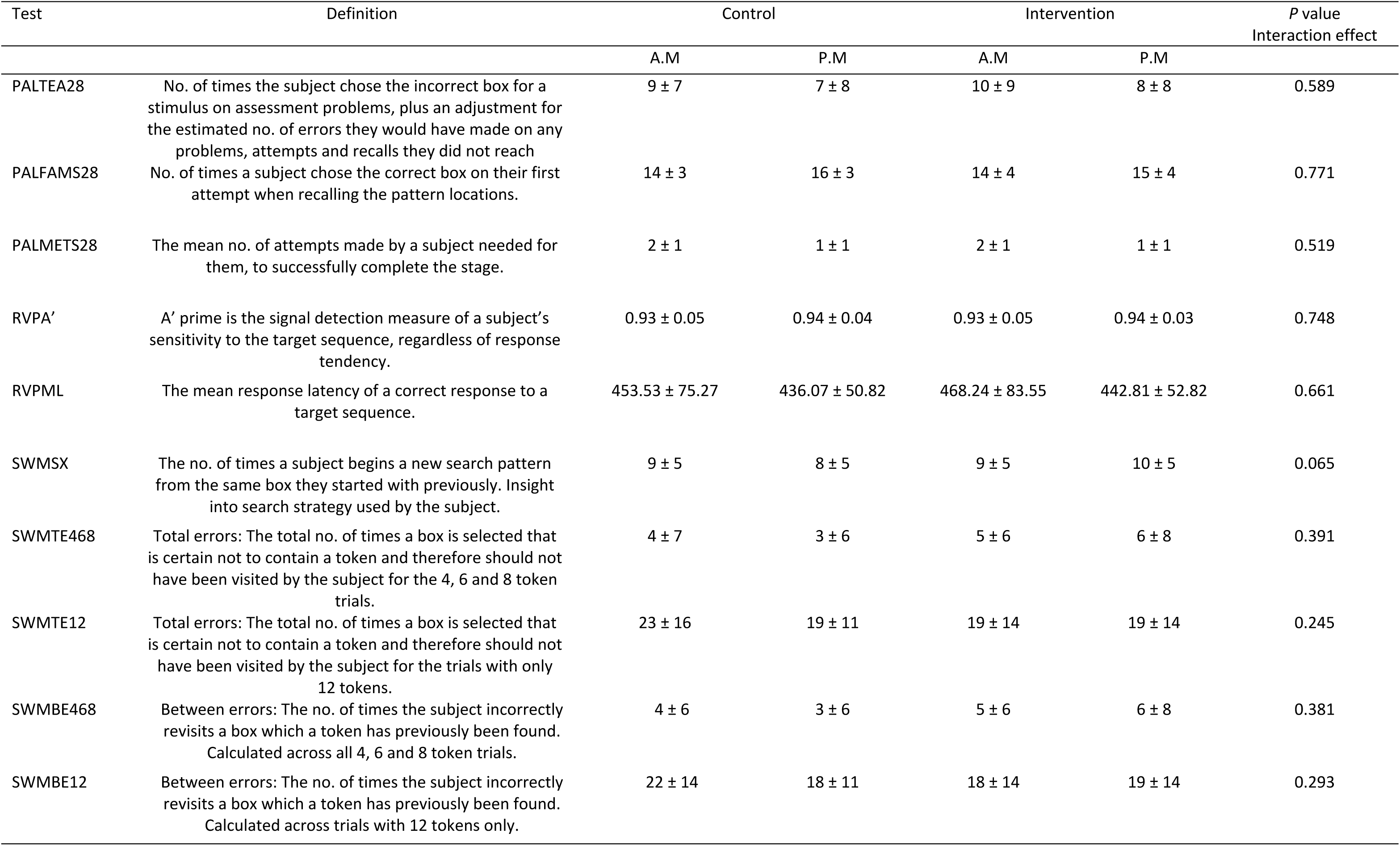
Mean ± SD results and *P* values for interaction effects in PAL, RVP and SWM cognitive assessment outputs.

| Test | Definition | Control |  | Intervention |  | *P* value<br>Interaction effect |
| --- | --- | --- | --- | --- | --- | --- |
|  |  | A.M | P.M | A.M | P.M |  |
| PALTEA28 | No. of times the subject chose the incorrect box for a stimulus on assessment problems, plus an adjustment for the estimated no. of errors they would have made on any problems, attempts and recalls they did not reach | 9 ± 7 | 7 ± 8 | 10 ± 9 | 8 ± 8 | 0.589 |
| PALFAMS28 | No. of times a subject chose the correct box on their first attempt when recalling the pattern locations. | 14 ± 3 | 16 ± 3 | 14 ± 4 | 15 ± 4 | 0.771 |
| PALMETS28 | The mean no. of attempts made by a subject needed for them, to successfully complete the stage. | 2 ± 1 | 1 ± 1 | 2 ± 1 | 1 ± 1 | 0.519 |
| RVPA' | A' prime is the signal detection measure of a subject's sensitivity to the target sequence, regardless of response tendency. | 0.93 ± 0.05 | 0.94 ± 0.04 | 0.93 ± 0.05 | 0.94 ± 0.03 | 0.748 |
| RVPML | The mean response latency of a correct response to a target sequence. | 453.53 ± 75.27 | 436.07 ± 50.82 | 468.24 ± 83.55 | 442.81 ± 52.82 | 0.661 |
| SWMSX | The no. of times a subject begins a new search pattern from the same box they started with previously. Insight into search strategy used by the subject. | 9 ± 5 | 8 ± 5 | 9 ± 5 | 10 ± 5 | 0.065 |
| SWMTE468 | Total errors: The total no. of times a box is selected that is certain not to contain a token and therefore should not have been visited by the subject for the 4, 6 and 8 token trials. | 4 ± 7 | 3 ± 6 | 5 ± 6 | 6 ± 8 | 0.391 |
| SWMTE12 | Total errors: The total no. of times a box is selected that is certain not to contain a token and therefore should not have been visited by the subject for the trials with only 12 tokens. | 23 ± 16 | 19 ± 11 | 19 ± 14 | 19 ± 14 | 0.245 |
| SWMBE468 | Between errors: The no. of times the subject incorrectly revisits a box which a token has previously been found. Calculated across all 4, 6 and 8 token trials. | 4 ± 6 | 3 ± 6 | 5 ± 6 | 6 ± 8 | 0.381 |
| SWMBE12 | Between errors: The no. of times the subject incorrectly revisits a box which a token has previously been found. Calculated across trials with 12 tokens only. | 22 ± 14 | 18 ± 11 | 18 ± 14 | 19 ± 14 | 0.293 |

## Discussion

This study investigated the effects of morning RT on afternoon physical and cognitive performance within male RU players. Results demonstrated that afternoon physical performance was significantly improved following morning RT. However, no significant improvements in cognitive performance were discovered despite RVPML being influenced by time.

Relative to CON, completing a combination of lower body strength and ballistic exercise enhanced CMJ flight time (0.385 ms ± 0.053, 2%) and 30 m fly and 40 m sprinting performance (-0.08 ± 0.13 s, 8%, -0.12 ± 0.17 s, 12%, respectively). These improvements in afternoon physical performance extends recent recommendations that morning RT of low volume, high-intensity RT can improve neuromuscular performance six hours later [8]. It has been reported physical performance improvements following both high intensity RT (> 80% 1RM) and ballistic exercise performed with maximal intent at lower intensities ( ≤ 45% 1RM) [30]. This research has not, however, combined both methods within the same priming session. Our findings demonstrate that combining both can enhance afternoon physical performance within male team sport athletes - specifically the lower body. It would be useful for future research to investigate whether the effects extend to upper body performance, are sex specific and the effect on amateur and professional athletes. Notably, the priming response to a stimulus incorporating the barbell squat jump exercise has not previously been assessed within the recommended time frame of 5-6 hours [30, 31]. Our findings advocate for the use of this exercise within a morning RT primer and may well be more accepted by elite athletes on competition day.

In the case of sprinting, our results provide further insight to the effects of morning RT on afternoon sprinting performance, where there is currently a scarcity of research. Cook et al [6] reported a decrease in 40 m sprint times (-0.02 - 0.08 s) whereby subjects completed a three-RM in the barbell back squat and bench press exercises. In contrast, the current study demonstrates that significant sprint improvements may be found with sub-maximal intensity which may be more feasible on day of competition. Additionally, our findings of improved sprint performance following a lower body RT session despite recent suggestions that the ergogenic effects of priming are movement specific [9]. Sprinting itself has been researched as a method of priming and has demonstrated positive results [9]. Despite its practicality due to no equipment required, athletes may not wish to maximally sprint on the morning of competition. The results of the current study extend the agreement of previous literature, demonstrating that morning RT can improve afternoon sprinting performance and may display a greater ergogenic effect than sprinting alone [7, 15].

The literature is currently unclear on the effects of morning RT on cognitive performance, with previous literature investigating reaction time only [7, 15]. In the current study no significant improvements in cognitive performance, specifically reaction time, sustained attention and working memory were found, supporting previous findings within elite rugby players [7]. Although non-significant, lower response latencies were found in the afternoon following INT compared to CON (-5.56% vs. -3.75%, respectively) in the RVP assessment in the current study. Despite differing assessments, previous evidence in professional cricketers has shown significant improvements in reaction time following morning RT [15]. However, the mechanism behind these results is currently unknown. Nutt et al [15] as with the current study did not undertake hormonal assessment. As with the physical performance improvement mechanisms, the hormone testosterone may play a role in acutely enhancing cognitive performance. On the contrary, the findings of Russell et al [7] reject the notion of testosterone’s role on cognitive performance, at least acutely. Alternatively, multiple cognitive abilities have been shown to fluctuate throughout the day, with different abilities peaking at different time-points [32, 33]. For example, one study reported improvements in dart-throwing accuracy as the day progressed, with best accuracy recorded in the evening (19:00 h) compared to the afternoon (15:00 h) and better than the morning (07:00 h) [34]. Furthermore, the lack ecological validity of the cognitive assessments used is a noteworthy limitation of the current study. Future research investigating sporting skill performance such as passing accuracy under time pressure within RU may provide a more ecologically valid insight to the effects of a morning RT on afternoon cognitive performance. Additionally, it is currently unknown whether a sporting skill-based cognitive primer completed in the morning may improve afternoon performance. In the case of RU, this may involve the completion of a three versus two attack and defence drill that have various cognitive requirements such as decision making and attention.

The mechanisms behind our observed results cannot be fully determined from the present study, however previous researchers have discovered that morning RT can have positive effects offsetting the diurnal decline of testosterone throughout the day, leading to enhanced physical and potentially cognitive afternoon performance [6, 15]. The lack of physiological and hormonal analysis is a limitation of the current study which must be noted. The physical improvements reported despite no cognitive improvement suggest other mechanisms may play a significant role. For example, muscle and body temperature also demonstrate diurnal variations throughout the day and correlate with physical performance indices [35, 36].

Furthermore, it has been suggested that neuromuscular responses may play a key role in the priming response [8]. This theory would support the findings within the current study; however, further research is warranted. Notably, despite asking subjects to maintain habitual nutritional habits during assessment days in the current study, the timing and macronutrient composition of food during the passive recovery phase may have had detrimental effects on their afternoon cognitive performance [37]. Future research may look to control for creatine and caffeine alongside timing and type of food ingestion due to their effects on cognitive performance [38, 39]. Due to the current mixed findings further research investigating how cognitive performance may be acutely enhanced to support athletes on competition day is warranted.

### Practical application

Morning RT consisting of heavy strength (barbell back squat, three sets of three repetitions at 85% RM) and ballistic (barbell squat jump, five sets of three repetitions at 40% RM) exercise can improve indices of afternoon physical performance (CMJ and sprinting performance) when commenced 5.5-hours after the priming stimulus. Applied practitioners may wish to prescribe low volume, high intensity, and ballistic lower body resistance exercise on the morning of competition to improve afternoon sprinting and jumping performance within male team sport athletes. However, the effects of cognitive performance remain unclear.

### Conclusions

We investigated the effects of a morning RT priming session on afternoon physical and cognitive performance. Our results demonstrate that afternoon physical performance can be improved after a 5.5-hour time interval that followed low volume, lower body morning RT in male rugby union players. Future studies should determine whether these results can be extended to other team sports and female athletes. Conversely, our results did not support significant improvements in afternoon cognitive performance.

## Supporting Information

S1 File. (Table A) Linear sprint test results. (Table B) Countermovement jumps test results. (Table C) Cognitive assessment test results.

## Acknowledgements

No acknowledgments to declare.

## Author Contributions

**Conceptualisation:** J. Knight, L. Kilduff, M. Russell

**Data curation:** J. Knight, D. Cunningham

**Formal analysis**: J. Knight, L. Kilduff, M. Russell

**Investigation:** J. Knight

**Methodology:** J. Knight, L. Kilduff, M. Russell

**Project administration:** J. Knight

**Supervision:** L. Kilduff, M. Russell

**Writing – original draft:** J. Knight, L. Kilduff, M. Russell

**Writing – review &editing:** D. Cunningham, C. Cook, M. Waldron

## Notes

### Competing Interest Statement

The authors have declared no competing interest.

## References

1. Cunningham DJ, Shearer DA, Drawer S, Pollard B, Cook CJ, Bennett M, et al. Relationships between physical qualities and key performance indicators during match-play in senior international rugby union players. Rogan S, editor. PLOS ONE. 2018 Sep 12;13(9). 10.1371/journal.pone.0202811. PMID: 30208066.

2. Bennett M, Bezodis N, Shearer DA, Locke D, Kilduff LP. Descriptive conversion of performance indicators in rugby union. Journal of Science and Medicine in Sport. 2019 Mar;22(3):330–4. 10.1016/j.jsams.2018.08.008. PMID 30146476.

3. Waldron M, Worsfold PR, Twist C, Lamb K. The relationship between physical abilities, ball-carrying and tackling among elite youth rugby league players. Journal of Sports Sciences. 2013 Sep 27;32(6):542–9.

4. Kruger A, Du Plooy K, Kruger P. Thinking differently about rugby performance: The relationship between cognitive functioning and on-field performance. African Journal for Physical Health Education, Recreation and Dance. 2018 Mar 1;24.

5. Kilduff LP, Pyne DB, Cook CJ. Performance Science Domains: Contemporary Strategies for Teams Preparing for the Rugby World Cup. International Journal of Sports Physiology and Performance [Internet]. 2023 Aug 18;18(9):1085–8. 10.1123/ijspp.2023-0179. PMID: 37573027.

6. Cook CJ, Kilduff LP, Crewther BT, Beaven M, West DJ. Morning based strength training improves afternoon physical performance in rugby union players. Journal of Science and Medicine in Sport. 2014;17(3):317–21. 10.1016/j.jsams.2013.04.016. PMID: 23707139.

7. Russell M, King A, Bracken RM, Cook CJ, Giroud T, Kilduff LP. A comparison of different modes of morning priming exercise on afternoon performance. International Journal of Sports Physiology and Performance. 2016 Sep;11(6):763–7. 10.1123/ijspp.2015-0508. PMID: 26658460.

8. Harrison PW, James LP, McGuigan MR, Jenkins DG, Kelly VG. Resistance Priming to Enhance Neuromuscular Performance in Sport: Evidence, Potential Mechanisms and Directions for Future Research. Vol. 49, Sports Medicine. Springer International Publishing; 2019. p. 1499–514. 10.1007/s40279-019-01136-3. PMID: 31203499.

9. Mason B, McKune A, Pumpa K, Ball N. The Use of Acute Exercise Interventions as Game Day Priming Strategies to Improve Physical Performance and Athlete Readiness in Team-Sport Athletes: A Systematic Review. Vol. 50, Sports Medicine. Springer Science and Business Media Deutschland GmbH; 2020. p. 1943–62. 10.1007/s40279-020-01329-1. PMID: 32779102.

10. MacDonald LA, Minahan CL. Indices of cognitive function measured in rugby union players using a computer-based test battery. Journal of Sports Sciences. 2016 Sep;34(17):1669–74. 10.1080/02640414.2015.1132003. PMID: 26756946.

11. Furley PA, Memmert D. Working memory capacity as controlled attention in tactical decision making. Journal of Sport and Exercise Psychology. 2012;34(3):322–44. 10.1123/jsep.34.3.322. PMID: 22691397.

12. Scharfen HE, Memmert D. The relationship between cognitive functions and sport-specific motor skills in elite youth soccer players. Frontiers in Psychology. 2019;10(APR). 10.3389/fpsyg.2019.00817. PMID: 31105611.

13. Huang TY, Chen FT, Li RH, Hillman CH, Cline TL, Chu CH, et al. Effects of Acute Resistance Exercise on Executive Function: A Systematic Review of the Moderating Role of Intensity and Executive Function Domain. Vol. 8, Sports Medicine - Open. Springer Science and Business Media Deutschland GmbH; 2022. 10.1186/s40798-022-00527-7. PMID: 36480075.

14. Tsuk S, Netz Y, Dunsky A, Zeev A, Carasso R, Dwolatzky T, et al. The acute effect of exercise on executive function and attention: Resistance versus aerobic exercise. Advances in Cognitive Psychology. 2019 Sep;15(3):208–15. 10.5709/acp-0269-7. PMID: 32161629.

15. Nutt F, Hills SP, Russell M, Waldron M, Scott P, Norris J, et al. Morning resistance exercise and cricket-specific repeated sprinting each improve indices of afternoon physical and cognitive performance in professional male cricketers. Journal of Science and Medicine in Sport. 2022 Feb;25(2):162–6. 10.1016/j.jsams.2021.08.017. PMID: 34535402.

16. Trumble BC, Stieglitz J, Thompson ME, Fuerstenberg E, Kaplan H, Gurven M. Testosterone and male cognitive performance in Tsimane forager-horticulturalists. American Journal of Human Biology. 2015 Jul;27(4):582–6. 10.1002/ajhb.22665. PMID: 25429990.

17. Kraemer WJ, Ratamess NA. Hormonal Responses and Adaptations to Resistance Exercise and Training. Vol. 35, Sports Med. 2005. p. 339–61.

18. Villarreal ESS de, González-Badillo JJ, Izquierdo M. Optimal warm-up stimuli of muscle activation to enhance short and long-term acute jumping performance. European Journal of Applied Physiology. 2007 Jul;100(4):393–401. 10.1007/s00421-007-0440-9. PMID: 17394010.

19. Mason BRJ, Argus CK, Norcott B, Ball NB. Resistance Training Priming Activity Improves Upper-Body Power Output in Rugby Players. Journal of Strength and Conditioning Research. 2017 Apr;31(4):913–20. PMID: 27386962.

20. Teo W, Newton MJ, McGuigan MR. Circadian rhythms in exercise performance: implications for hormonal and muscular adaptation. Journal of Sports Science & Medicine [Internet]. 2011 Dec 1;10(4):600–6. Available from: https://pubmed.ncbi.nlm.nih.gov/24149547/. PMID: 24149547.

21. Karlsen RH, Karr JE, Saksvik SB, Lundervold AJ, Hjemdal O, Olsen A, et al. Examining 3-month test-retest reliability and reliable change using the Cambridge Neuropsychological Test Automated Battery. Applied Neuropsychology:Adult. 2022;29(2):146–54. 10.1080/23279095.2020.1722126. PMID: 32083946.

22. Gonçalves MM, Pinho MS, Simões MR. Test–retest reliability analysis of the Cambridge neuropsychological automated tests for the assessment of dementia in older people living in retirement homes. Applied Neuropsychology:Adult. 2016 Jul;23(4):251–63. 10.1080/23279095.2015.1053889. PMID: 26574661.

23. Gonçalves MM, Pinho MS, Simões MR. Construct and concurrent validity of the Cambridge neuropsychological automated tests in Portuguese older adults without neuropsychiatric diagnoses and with Alzheimer’s disease dementia. Aging, Neuropsychology, and Cognition. 2018 Mar;25(2):290–317. 10.1080/13825585.2017.1294651. PMID: 2863207.

24. Smith PJ, Need AC, Cirulli ET, Chiba-Falek O, Attix DK. A comparison of the Cambridge Automated Neuropsychological Test Battery (CANTAB) with “traditional” neuropsychological testing instruments. Journal of Clinical and Experimental Neuropsychology. 2013 Feb 27;35(3):319–28. 10.1080/13803395.2013.771618. PMID: 23444947.

25. Torgersen J, Flaatten H, Engelsen BA, Gramstad A. Clinical Validation of Cambridge Neuropsychological Test Automated Battery in a Norwegian Epilepsy Population. Journal of Behavioral and Brain Science. 2012;02(01):108–16. 10.4236/jbbs.2012.21013

26. Backx R, Skirrow C, Dente P, Barnett JH, Cormack FK. Comparing Web-Based and Lab-Based Cognitive Assessment Using the Cambridge Neuropsychological Test Automated Battery: A Within-Subjects Counterbalanced Study. Journal of Medical Internet Research. 2020 Aug 4;22(8):e16792. 10.2196/16792. PMID: 32749999.

27. Owen N, Watkins J, Kilduff L, Bevan H, Bennett M. Development of a Criterion Method to Determine Peak Mechanical Power Output in a Countermovement Jump. The Journal of Strength and Conditioning Research. 2014 Jun 1;28:1552–8. 10.1519/jsc.0000000000000311. PMID: 24276298.

28. Cohen J. Statistical power analysis. Current Directions in Psychological Science. 1992 Jun;1(3):98–101.

29. Holmberg PM, Harrison PW, Jenkins DG, Kelly VG. Factors Modulating the Priming Response to Resistance and Stretch-Shortening Cycle Exercise Stimuli. Strength and Conditioning Journal. 2023; 45(2): 188–206. 10.1519/ssc.0000000000000728.

30. Tsoukos A, Veligekas P, Brown LE, Terzis G, Bogdanis GC. Delayed Effects of a Low-Volume, Power-Type Resistance Exercise Session on Explosive Performance. Journal of Strength and Conditioning Research. 2018 Mar;32(3):643–50. 10.1519/jsc.0000000000001812. PMID: 28291764.

31. Nishioka T, Okada J. Influence of Strength Level on Performance Enhancement Using Resistance Priming. Journal of Strength and Conditioning Research. 2021 Oct 27;36(1):37–46. 10.1519/jsc.0000000000004169. PMID: 34711771.

32. Carrier J, Monk TH. CIRCADIAN RHYTHMS OF PERFORMANCE: NEW TRENDS. Chronobiology International. 2000 Jan;17(6):719–32. 10.1081/cbi-100102108. PMID: 11128289.

33. Van Dongen HPA, Dinges DF. Sleep, Circadian Rhythms, and Psychomotor Vigilance. Clinics in Sports Medicine. 2005 Apr;24(2):237–49. 10.1016/j.csm.2004.12.007. PMID: 15892921.

34. Edwards B, Waterhouse J, Atkinson G, Reilly T. Effects of time of day and distance upon accuracy and consistency of throwing darts. Journal of Sports Sciences. 2007; 25(13), 1531–1538. 10.1080/02640410701244975. PMID: 17852679.

35. Pullinger SA, Brocklehurst EL, Iveson RP, Burniston JG, Doran DA, Waterhouse JM, Edwards BJ. Is there a diurnal variation in repeated sprint ability on a non-motorised treadmill? Chronobiology International. 2014; 31(3), 421–432. 10.3109/07420528.2013.865643. PMID: 24328815.

36. Pullinger SA, Oksa J, Clark LF, Guyatt, JWF, Newlove A, Burniston JG, et al. Diurnal variation in repeated sprint performance cannot be offset when rectal and muscle temperatures are at optimal levels (38.5°C). Chronobiology International. 2018; 35(8), 1054–1065. 10.1080/07420528.2018.1454938. PMID: 29566344.

37. Mahoney C, Taylor H, Kanarek R. 6 The Acute Effects of Meals on Cognitive Performance. Nutritional Neuroscience. 2005 Mar 18; 73–91. https://psycnet.apa.org/doi/10.1201/9780203564554.ch6

38. Walczak K, Krasnoborska J, Samojedny S, Superson M, Szmyt K, Szymańska K, Wilk-Trytko K. Effect of creatine supplementation on cognitive function and mood. J Educ Health Sport [Internet]. 2024 Jun. 3; 73:51712.

39. McLellan TM, Caldwell JA, Lieberman HR. A review of caffeine’s effects on cognitive, physical and occupational performance. Neuroscience & Biobehavioral Reviews [Internet]. 2016 Dec;71(1):294–312. Available from: https://www.sciencedirect.com/science/article/pii/S0149763416300690

